# Identifying structural variants using linked-read sequencing data

**DOI:** 10.1101/190454

**Authors:** Rebecca Elyanow, Hsin-Ta Wu, Benjamin J. Raphael

**Affiliations:** Department of Computer Science, Princeton University, Princeton, NJ 08540.; Center for Computational Molecular Biology, Brown University, Providence, RI 02912.

## Abstract

Structural variation, including large deletions, duplications, inversions, translocations, and other rearrangements, is common in human and cancer genomes. A number of methods have been developed to identify structural variants from Illumina short-read sequencing data. However, reliable identification of structural variants remains challenging because many variants have breakpoints in repetitive regions of the genome and thus are difficult to identify with short reads. The recently developed linked-read sequencing technology from 10X Genomics combines a novel barcoding strategy with Illumina sequencing. This technology labels all reads that originate from a small number (~5-10) DNA molecules ~50Kbp in length with the same molecular barcode. These barcoded reads contain long-range sequence information that is advantageous for identification of structural variants. We present Novel Adjacency Identification with Barcoded Reads (NAIBR), an algorithm to identify structural variants in linked-read sequencing data. NAIBR predicts novel adjacencies in a individual genome resulting from structural variants using a probabilistic model that combines multiple signals in barcoded reads. We show that NAIBR outperforms several existing methods for structural variant identification – including two recent methods that also analyze linked-reads – on simulated sequencing data and 10X whole-genome sequencing data from the NA12878 human genome and the HCC1954 breast cancer cell line. Several of the novel somatic structural variants identified in HCC1954 overlap known cancer genes.

## 1 Introduction

Recent whole genome sequencing (WGS) analysis of human genomes has shown that *structural variation*, including insertions, deletions, duplications, and rearrangements of genomic segments greater than 50bp, are a key component of human variation (Sudmant *et al*., 2015). Collectively, structural variants affect a larger portion of the human genome than single nucleotide variants (Pang *et al*., 2010). Inherited germline structural variants have been implicated in several diseases including Crohn’s disease, rheumatoid arthritis, Type I diabetes, and autism (Mccarroll *et al*., 2009; Weisenfeld *et al*., 2016; Sebat *et al*., 2007). In addition, somatic structural variants are common in cancer genomes (Chittenden *et al*., 2008; Rausch *et al*., 2013). These include deletions of tumor suppressor genes and amplifications of oncogenes which can promote aggressive cell growth and drive the development of cancer. Cancer genomes can also undergo dramatic rearrangement events such as chromothripsis, the shattering and random repair of chromosomes in a single catastrophic event (Stephens *et al*., 2011), or chromoplexy (Baca *et al*., 2013), both of which result in a large number of complex structural variants in a cancer genome.

The identification of structural variants from high-throughput DNA sequencing data is generally more challenging than the identification of single nucleotide variants. This difficulty is primarily a result of the the fact that many structural variants are significantly longer than the DNA sequence reads produced by current (second generation) DNA sequencing technologies, whose read lengths are ~300-500 nucleotides. In addition, such reads are too short for *de novo* genome assembly. Thus structural variants are inferred from atypical, or aberrant, alignments of reads to a reference genome.

Numerous methods have been developed over the past several years to identify different types of structural variants from read alignments. Each of these methods use some combination of three signals that can be extracted from read alignments: *discordant read-pairs, split-reads*, and *read depth* (Figure S12a). A discordant read-pair is a pair of reads from the same fragment/insert whose alignments to the reference genome have distance and/or orientation that differ from expected if the entire fragment was contiguous on the reference genome. A split read is a read with no contiguous alignment to the reference genome, but rather with at least two partial alignments to the reference. (In practice, only a single partial, or clipped, alignment may be reported by the read alignment software.) Discordant read pairs and split reads are signatures of a *novel adjacency* in the individual genome; that is, two intervals that are non-adjacent in the reference genome are adjacent in an individual genome. Methods that rely on discordant paired reads and/or split reads include BreakDancer (Chen *et al*., 2009), GASV (Sindi *et al*., 2009a), VariationHunter (Hormozdiari *et al*., 2010), Pindel (Ye *et al*., 2009), DELLY (Rausch *et al*., 2012), and LUMPY (Layer *et al*., 2014); many others are reviewed in Tattini *et al*. (2015). Read depth is the (normalized) number of reads that map to a particular region of the genome. Read depth can be used to identify copy-number aberrations, such as deletions and duplications, by identifying regions of unexpectedly low or high coverage in the genome. Examples of read depth methods include BIC-Seq (Xi *et al*., 2010), and CNVnator (Abyzov *et al*., 2011). In addition, methods such as GASVPro (Sindi *et al*., 2012) and SV-Bay (Iakovishina *et al*., 2016) combine signals from discordant read-pairs and read depth signals to identify structural variants. Local assembly approaches such as SvABA (Wala *et al*., 2017), and novoBreak (Chong *et al*., 2017) achieve nucleotide level resolution of novel adjacencies. However, local assembly approaches require the identification of candidate regions for assembly using the signals described above; thus, local assembly generally increases specificity more than sensitivity. In addition, assembly-based approaches requires high coverage and is typically more time consuming than read-pair or read depth based methods (Alkan *et al*., 2011).

The fundamental limitations in structural variant detection and whole-genome assembly are the fact that the human genome is diploid and highly repetitive (Mak *et al*., 2016). Short reads have low signal to assign variants to haplotypes and to identify structural variants whose breakpoints lie in repetitive sequences. While algorithms can help extract information from this low signal, longer reads provide stronger signal. New *3rd-generation* sequencing technologies developed by Pacific Biosciences and Oxford Nanopore produce much longer reads (exceeding 10Kbp). However, these technologies are practically limited by their high per-base error rate and high cost compared to Illumina short-read sequencing. Additionally, structural variants that are larger than the average read size or that fall in repetitive regions still remain difficult to identify using these technologies (Weischenfeldt *et al*., 2013). An alternative sequencing technology called *linked-read sequencing* was recently developed by 10X Genomics, and commercialized in their Chromium platform. In this technology, long DNA molecules, 50 – 100Kbp in size are partitioned into one of several million droplets using microfluidics. Each droplet contains a small number (^~^10) of molecules (Weisenfeld *et al*., 2016). The molecules in each droplet are sheared into smaller fragments and labeled with a 16bp molecular barcode that is unique to each droplet. The fragments are then amplified and sequenced using Illumina paired-end sequencing protocol (Figure 1). 10X Genomics’ linked-read technology thus provides both the low error rate and low cost of Illumina sequencing as well as long-range sequencing information provided by 3rd-generation sequencing technologies (Figure S12b). The technology is similar in some respects to the strobe sequencing technology that was prototyped by Pacific BioSciences but never commercially released (Ritz *et al*., 2010, 2014).

**Figure 1:**
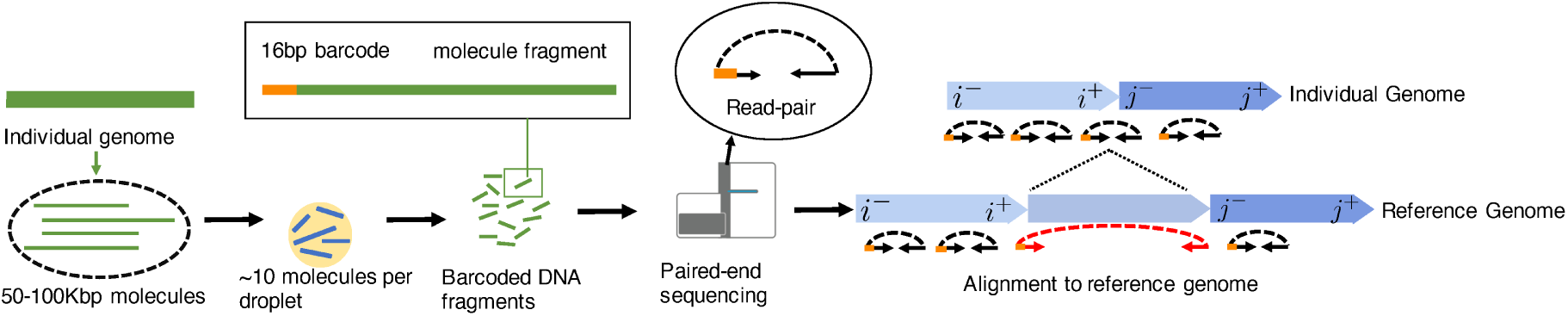
(Left) Linked-read sequencing with the 10X Genomics Chromium platform begins by fragmenting the individual genome into large DNA molecules, which are isolated into individuals beads that contain several large molecules and sequencing reagents. Within the bead, molecules are sheared into smaller fragments (500 bp) and labeled with a 16bp barcode indicating its bead of origin. Illumina paired-read sequencing of each fragment results in barcoded paired-end reads. (Right) Alignment of read-pairs to a reference genome results in concordant reads (black) and discordant reads (red). Discordant reads indicate candidate novel adjacencies that are a result of structural variants that distinguish individual genomes from the reference genome.

Here, we introduce Novel Adjacency Identification with Barcoded Reads (NAIBR, pronounced “neighbor”), a method that identifies novel adjacencies resulting from structural variants in an individual genome from linked-read sequencing data. NAIBR combines a novel split-read type signal from linked-reads with traditional signals of structural variants in the underlying paired-reads in the data. We demonstrate that NAIBR outperforms existing structural variant detection algorithms – both paired-read methods and two recently developed methods (Zheng *et al*., 2016; Spies *et al*., 2016) for linked-reads – using simulated and real linked-read sequencing data. NAIBR also leverages haplotype phasing information from linked-reads, improving the detection of heterozygous structural variants.

## 2 Methods

Consider two genomes, a *reference genome* and an *individual genome*, each represented by an interval, *G*= [1,*n*] and *G′*= [1,*n′*] respectively. We let [*i*^‒^, *i*^+^] denote an interval in the genome, where *i*^‒^ and *i*^+^ indicate the start and end of the interval respectively. We consider a structural variant to be any difference between an individual genome and a reference genome due to DNA breakage and repair, that results in the joining of two non-adjacent intervals [*i*^+^,*i*^‒^] and [*j*^+^, *j*^‒^] in the reference genome. The ends of these intervals may be joined in one of four orientations, and we indicate the four possible *novel adjacencies* in the individual genome by the pairs of joined ends: (*i*^+^, *j*^‒^),(*i*^+^, *j*^+^),(*i*^‒^, *j*^‒^), or (*i*^‒^, *j*^+^). Note that novel adjacencies are formed by most of the usual structural variants (segmental deletions/insertions, inversions, and translocations), with the notable exception of the deletion of a chromosome to the telomere.

NAIBR aims to identify such novel adjacencies using linked-read sequencing data. Our algorithm differs from previously published methods in that it incorporates signals from both paired-end reads and linked-reads into a unified model. Before defining the model, we will describe the signals we observe in paired-end and linked-read data and how these signals are combined to identify novel adjacencies arising from structural variants in an individual genome.

### 2.1 Paired-end sequencing data

In Illumina paired-end sequencing, chromosomes are sheared into small fragments and size selected such that most fragment lengths are within the interval [*l*_min_, *l*_max_]. Each fragment is sequenced from both ends from opposite DNA strands; thus one read will originate from the forward (+) strand and one from the reverse (-) strand. Paired reads are aligned to the reference genome. Each aligned read, *x*, is represented by a tuple *x* = (*l*_*x*_, *r*_*x*_, *o*_*x*_, *q*_*x*_), where *l*_*x*_ is the leftmost position of *x* in the reference genome, *r*_*x*_ is the rightmost position in the reference genome, *o*_*x*_ ∈ {+,-} is the orientation of *x*, and *q*_*x*_ is the mapping probability of *x*. We define a read-pair to be the ordered pair 〈*x, y*〉, where read *x* has the smaller starting coordinate. A read-pair 〈*x, y*〉 is *concordant* provided the distance between aligned reads *f* = *r*_*y*_ – *l*_*x*_ isbetween *l*_min_ and *l*_max_ and the orientations are *o*_*x*_ = +, *o*_*y*_ = –. Concordant reads are consistent with the fragment aligning contiguously to the reference genome with no rearrangement. A read pair 〈*x, y*〉 not satisfying this condition is *discordant*. Discordant read-pairs arise from either (1) errors in sequencing and/or alignment or (2) novel adjacencies in an individual genome with respect to the reference.

### 2.2 Linked-read sequencing data

The linked-read sequencing technology developed 10X Genomics adds a layer of structure to Illumina paired-end sequencing by tagging each DNA fragment with a barcode prior to sequencing (Figure 1). Barcoded read-pairs are aligned to the reference using a linked-read aware aligner called Lariat (Bishara *et al*., 2015). Lariat processes all read-pairs from a single barcode simultaneously, using the knowledge that reads originate from a small number of long molecules. Using this prior knowledge, it finds more unique mappings than other tools and can map to highly repetitive regions.

Read-pairs *p* = 〈*x, y*〉 originating from the same long molecule will each be tagged with the same barcode *b*_*p*_ ∈ ℕ. Because molecules are partitioned into droplets uniformly at random, the likelihood of assigning the same barcode to two molecules from nearby locations on the reference genome is low. Thus, we assume that read-pairs with the same barcode that map near each other on an individual genome are likely to have originated from the same long molecule. We partition such read-pairs into sets called *linked-reads*.

A linked-read is a set of concordant read-pairs that share the same barcode and have a maximum distance of *δ* from another read-pair in the linked-read. Each linked-read is assumed to have originated from a contiguous strand of DNA in the reference genome. The set of all concordant read-pairs in the genome is partitioned into linked-reads such that any two read-pairs *p* = 〈*x, y*〉 and *p′* = 〈*x*′, *y*′〉, where *l*_*x*_ < *l′*_*x*_, are both partitioned into the same linked-read if *b*_*p*_ = *b*_*p′*_, and *l*_*x′*_ – *r*_*y*_ ≤ *δ* (see Figure 2a).

**Figure 2:**
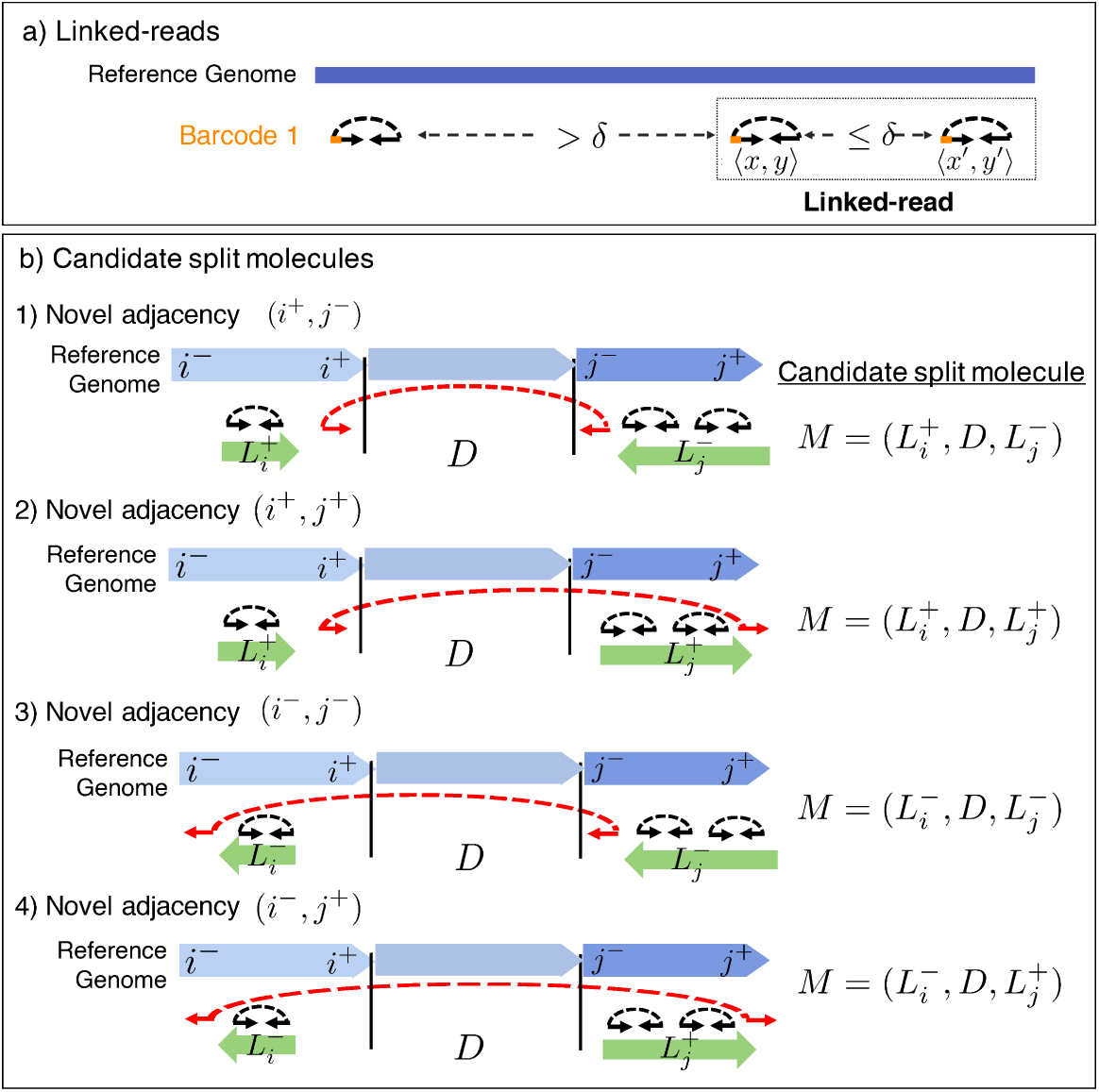
(a) A linked-read is defined by read-pairs separated by a distance ≤ *δ* on the reference genome. (b) Linked-reads *L*_*i*_ and *L*_*j*_ may have originated from one of 4 candidate split molecules, each supporting a novel adjacency with a different orientation. 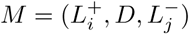 supports a novel adjacency (*i*^+^, *j*^‒^) and indicates that the end of linked-read *L*_*i*_ is adjacent to the start of linked-read *L*_*j*_ (the arrows points to the location of the novel adjacency).

Any pair of linked-reads sharing a barcode may have originated from a molecule that is split with respect to the reference genome due to the presence of a novel adjacency in an individual genome. We define a *candidate split molecule* as a tuple *M* = (*L*_1_, *D, L*_2_), where *L*_1_ and *L*_2_ are linked-reads and *D* is a set of discordant read pairs, satisfying the following conditions: (1) All read pairs in *L*_1_, *L*_2_, and *D* share the same barcode. (2) All discordant read pairs in *D* have the same orientations, and the distance between any two discordant read-pairs in *D* is at most *l*_max_. Formally, for read pairs *p* = 〈*x, y*〉,*p*′ = 〈*x, y*〉 ∈ *D*, |*x* – *x*′| < *l*_max_ and |*y* – *y*′| < *l*_max_. (3) The linked reads are located within *δ* of the discordant read-pairs in *D*, in the direction consistent with the orientation of *D*. To indicate the last condition, we assign an orientation to each linked read in *M*. For example, 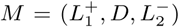 is a candidate split molecule provided: the *rightmost* position, 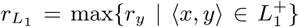, of *L*_1_ is within *δ* of discordant read-pairs in *D* and the *leftmost* position, 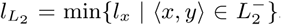, of *L*_2_ is within *δ* of discordant read-pairs in *D*. See Figure 2). Note that we also define a candidate split molecule in the case where *D* is the empty set. In this case, a candidate split molecule can be formed for any of the four possible orientations of linked-reads *L*_1_ and 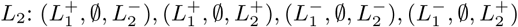.

We say that a candidate split molecule *M supports* a novel adjacency provided that the distances and orientations of the linked and discordant reads in *M* are consistent with the novel adjacency. For example, candidate split molecule 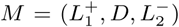 supports the novel adjacency (*i*^+^, *j*^‒^) provided: (1) The orientation of the candidate split molecule matches the orientation of the novel adjacency. (2) Each read-pair *p* = 〈*x, y*〉 ∈ *D*, is at most *l*_max_ from breakends *i* and *j*: *i* – *l*_max_ ≤ *r*_*x*_ ≤ *i* and *j* ≤ *l*_*y*_ ≤ *j* + *l*_max_. (3) Oriented linked-reads 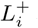 and 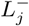 are at most a distance *δ* from positions *i* and *j* respectively: *i* – *δ* ≤ *r*_*L*_1__ ≤ *i* and *j* ≤ *l*_*L*_2__ ≤ *j* + *δ*.

According to the definitions above, for a given barcode and novel adjacency (*i*^+^, *j*^‒^), there is at most one candidate split molecule that supports this novel adjacency. We define the set *ℳ* to be the set of all candidate split molecules supporting a novel adjacency (*i*^+^, *j*^‒^). For ease of exposition, we will describe the model below for a novel adjacency of the form (*i*^+^, *j*^‒^), but the model may be applied to novel adjacencies with any of the four orientations.

### 2.3 Likelihood ratio score

We use a likelihood ratio score to evaluate the evidence support a novel adjacency. Given a potential novel adjacency (*i*^+^, *j*^‒^), let *A*_*i*^+^,*j*^‒^_ be the event of a novel adjacency (*i*^+^,*j*^‒^) in an individual genome and let event *A̅*_*i*^+^,*j*^‒^_, to be the absence of this novel adjacency. Let *ℳ* be the set of all candidate split molecules supporting novel adjacency (*i*^+^, *j*^‒^). We compare the likelihood *P*(*ℳ* | *A*_*i*+,*j*–_,*j-*) of *A*_*i*^+^,*j*^‒^_ and the likelihood *P*(*ℳ* | *A̅*_*i*^+,^*j*^‒^_ of *A̅*_*i*^+^,*j*^‒^_, given the set of observed candidate split molecules (Figure 3) using the log-likelihood ratio

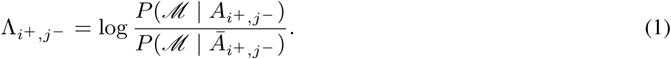

We report novel adjacencies with log-likelihood ratio, *A*_*i*^+^,*j*^‒^_ > *c* (selection of *c* is described in the Supplement) as the set of predicted novel adjacencies in the individual genome.

**Figure 3:**
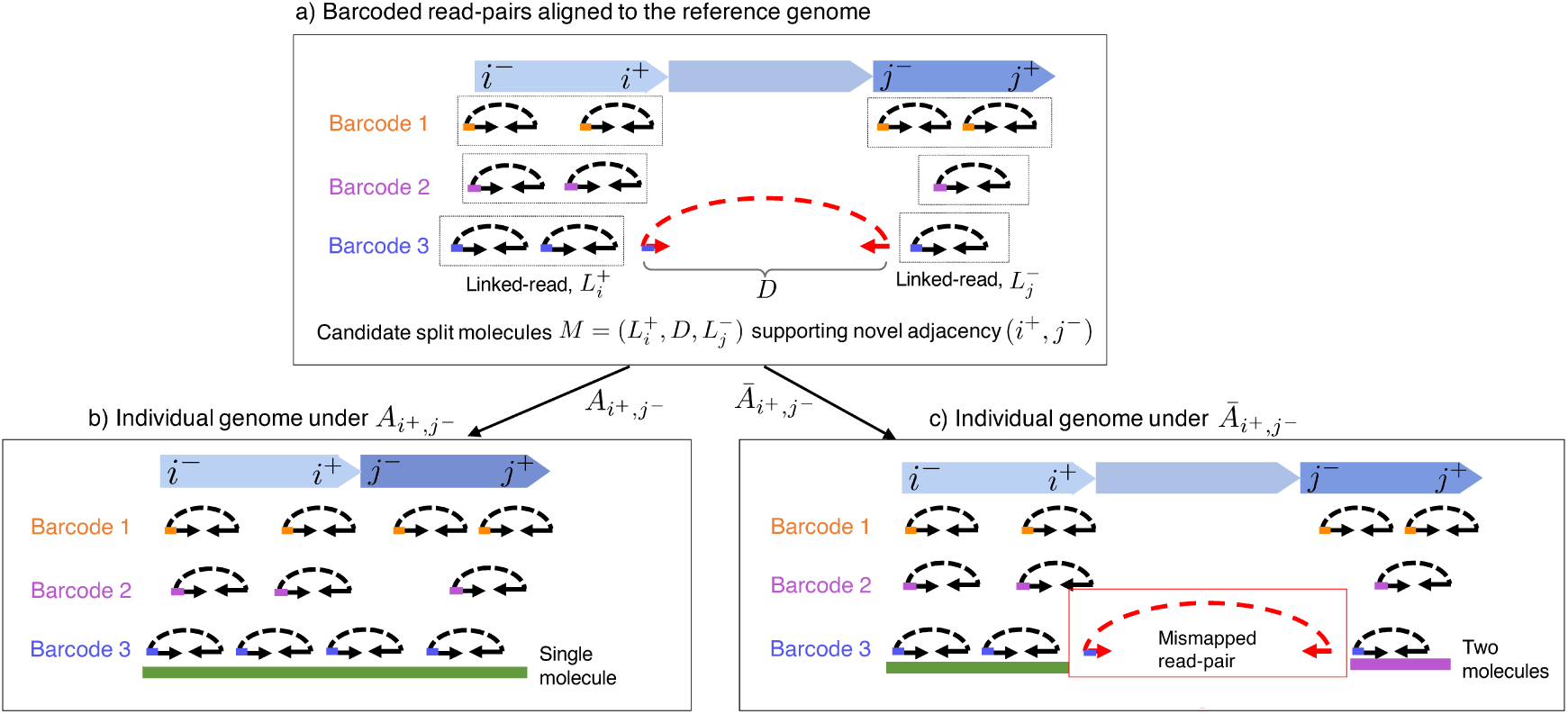
(a) A candidate split molecule for novel adjacency (*i*^+^, *j*^‒^) consists of a linked-read 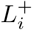 a linked-read 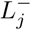, within a distance *δ* of position *j* and a set of discordant reads *D*. Barcoded reads aligned to the reference genome may originate from an individual genome that either contains a novel adjacency (*A*_*i*^+^,*j*^‒^_) or does not contain a novel adjacency (*A̅*_*i*^+^,*j*^‒^_). (b) Under the alternative hypothesis *A*_*i*^+^,*j*^‒^_ that *i*^+^ and *j*^‒^ are adjacent in an individual genome, reads in barcode 3 are close and are likely to have originated from a single molecule. (c) Under the null hypothesis *A̅*_*i*^+^,*j*^‒^_ that *i* and *j* are non-adjacent in an individual genome, reads in barcode 3 are separated by a large distance and are likely to have originated from two molecules. Barcode 2 contains a read that is discordant under *A̅*_*i*^+^,*j*^‒^_,*j–* and therefore assumed to be mismapped.

We now describe how we compute each of the terms in the log-likelihood ratio. First, we assume that the locations of molecules with different barcodes are independent. Combining this assumption with the requirement that candidate split molecules with different barcodes must originate from different molecules, we have that *P*(*ℳ* | *A*_*i*^+^ *j*^‒^_) and *P*(*ℳ* | *p* = *A*_*i*^+^,*j*^‒^_) are each the product of the individual probabilities of observing each candidate split molecule 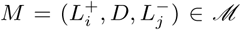. Next, a candidate split molecule *M* may have either originated from a single molecule that spans the interval [*i*^+^, *j*^‒^] or from two molecules, each of which does not span the interval [*i*^+^, *j*^‒^]. Let *E*_*M*_ be the event that candidate split molecule *M* originates from a single molecule and let 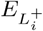 and 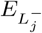 be the events that linked-reads 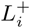 and 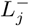 originate from unique molecules. Thus, the probability of observing candidate split molecule *M* is the probability of two disjoint events: that *M* originates from one molecule (*E*_*M*_), or that 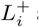 and 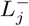 originate from two different molecules 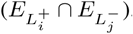. Thus, the log-likelihood ratio Ʌ_*i*^+^,*j*^‒^_ is calculated as follows,

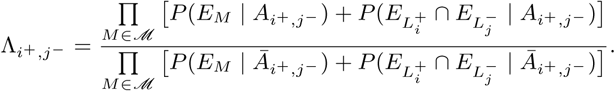

We now describe how we calculate the probabilities of *E*_*M*_ and 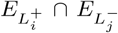 given the the events *A*_*i*^+^,*j*^‒^_ and *A̅*_*i*^+^,*j*^‒^_. Given a candidate split molecule 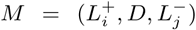) supporting a novel adjacency (*i*^+^,*j*^‒^), *P*(*E*_*M*_ | *A*_*i*+,*j*^ࢤ^_) is the probability that *M* was sequenced from a single molecule when *i*^+^ and *j*^‒^ are adjacent in the individual genome; correspondingly, *P*(*E*_*M*_ | *A̅*_*i*^+^,*j*^‒^_) is the probability that *M* was sequenced from a single molecule when *i*^+^ and *j*^‒^ are not adjacent in the individual genome. We model the sequencing of a molecule as the generation of mapped reads through three sequential processes: (1) molecule size selection, (2) molecule sequencing, and (3) read mapping. We assume that the fragmentation of chromosomes into long molecules and the sequencing of reads from a molecule are independent processes. This assumption of independence is reasonable because sequencing occurs after molecules are sheared into short fragments.

Thus, we model the probability *P*(*E*_*M*_ | *A*_*i*^+^,*j*^‒^_) as,

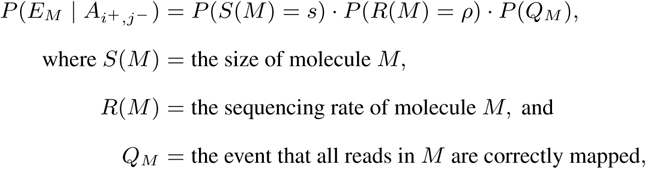

and the probability *P*(*E*_*M*_ | *A̅*_*i*^+^,*j*^‒^_) as,

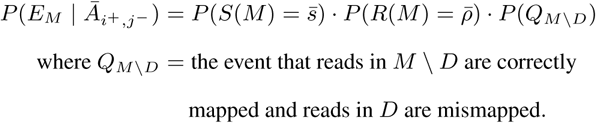

We model the size *S*(*M*) of a molecule *M* by a negative binomial distribution, *P*(*S*(*M*) = ·) = *NB*(·), with parameters estimated from the collection of all aligned linked reads. We assume that the vast majority of linked-reads in the data originate from molecules that are contiguous with respect to the reference genome. Formally, the size of a linked-read *L* is *r*_*L*_ – *l*_*L*_, where *r*_*L*_ = max{*r*_*y*_ | 〈*x, y*〉 ∈ *L*} and *l*_*L*_ = min{*l*_*x*_ | 〈*x, y*〉 ∈ *L*} are the rightmost and leftmost positions, respectively, of paired reads in *L*. This approximation tends to slightly underestimate the true length of a molecule due to missing reads at the ends of the molecule. Under *A*_*i*^+^,*j*^‒^_, the size of candidate split molecule *M* is approximately *s* = (*i* – *l*_*L*_*i*__) + (*r*_*L*_*j*__ – *j*), the sum of the portions of the molecule aligning to the left and right of the novel adjacency. Similarly, under *A̅*_*i*^+^,*j*^‒^_, the size of *M* is approximated by *s̅* = *r*_*L*_*i*__ – *l*_*L*_*i*__, the distance between the leftmost position in *L*_*i*_ and the rightmost position in *L*_*j*_. For the case where *i*^+^ and *j*^‒^ are on different chromosomes, *P*(*S*(*M*) = *s̅*) = 0.

We model the sequencing rate *R*(*M*) of a molecule *M* by a Gamma distribution, *P*(*R*(*M*) = ·) = Γ(·). The negative binomial and Gamma distributions provide a good fit to the empirical distributions of molecule size and sequencing rate respectively (Figure S2), but other distributions can be used (Table S1). The sequencing rate of a candidate split molecule *M* under *A̅*_*i*^+^,*j*^‒^_ is approximated by 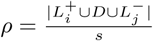, the number of reads sequenced from *M* normalized by the size *s* of *M*. The sequencing rate of *M* under *A̅*_*i*^+^,*j*^‒^_ is approximated by 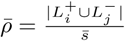. Under *A̅*_*i*^+^,*j*^‒^_, we exclude *D* from the set of reads sequenced from *M* because these reads are assumed to be mismapped.

We now define the probabilities *P*(*Q*_*M*_) and *P*(*Q*_*M*\*D*_) of mapping reads to the reference genome. Under *A*_*i*^+^,*j*^‒^_, all reads in *M* are correctly mapped, an event with probability

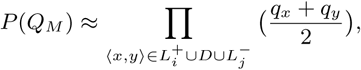

where *q*_*x*_ and *q*_*y*_ are the mapping probabilities of reads *x* and *y* obtained from the alignment software. We chose to approximate the mapping probability of the read-pair *p* = 〈*x, y*〉 to be the average of the mapping probabilities of each read, which we found to perform well on data. Under *A̅*_*i*^+^,*j*^‒^_, concordant reads in 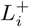 and 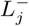 are correctly mapped and discordant reads in *D* are mismapped, an event with probability,

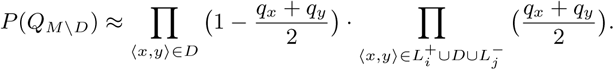

We calculate the probability 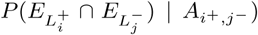 that 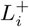 and 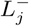 were sequenced from two different molecules as the product of the probabilities that 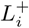 and 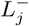 were sequenced from independent molecules with corresponding sizes *s*_*i*_, *s*_*j*_ and sequencing rates *p*_*i*_, *p*_*j*_, and that the reads from the two molecules were then properly mapped to the reference genome, the latter event denoted by event *Q*_*M*_. Formally,

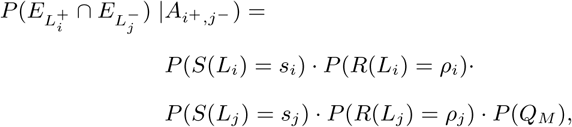

where *s*_*k*_ = *r*_*L*_*k*__ – *l*_*L*_*k*__ and 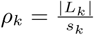.

Under *A̅*_*i*^+^,*j*^‒^_, the probabilities of molecule size selection and sequencing remain the same. However any discordant reads in *D* will be mismapped under *A̅*_*i*^+^,*j*^‒^_, resulting in the last term being the probability *P*(*Q*_*M*\*D*_).

### 2.4 Incorporating haplotype phase

10X Genomics provides phasing as part of its alignment pipeline, using linked-reads to phase SNPs into large phase blocks. Each position *i* in the reference genome is assigned a phase block *m*_*i*_ and any SNP within phase block *m*_*i*_ will be assigned to either haplotype 1 or haplotype 2. Zheng *et al*. (2016) report that the current software can phase > 95% of SNPs to a phase blocks of size > 0.5Mb, with an average error rate of 0.03%. Let *m*_*x*_ ∈ ℕ denote the phase block of a read *x*, aligned to the reference genome, and let *h*_*x*_ ∈ {1, 2} denote the haplotype of read *x*. We define *ℳ*^*α*,*β*^ ⊆ *ℳ* be the subset of candidate split molecules 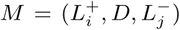 such that the haplotype *h*_*L*_*i*__ of linked-read *L*_*i*_ is *α*, the haplotype *h*_*L*_*j*__ of linked-read *L*_*j*_ is *β*, and for all read pairs 〈*x, y*〉 in *D, h*_*x*_ = *α* and *h*_*y*_ = *β*.

On the linked-read datasets described in the Results below, we found that > 99% of linked-reads (*δ* = 10Kbp) contain read-pairs that are assigned to a unique phase block and haplotype. We omit the small number of linked-reads containing read-pairs from multiple haplotypes or phase blocks from further analysis.

We define a haplotype-specific log-likelihood ratio,

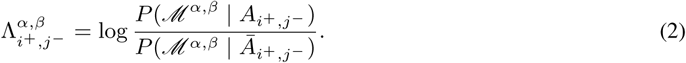

Each novel adjacency in a diploid genome is the result a heterozygous structural variant on haplotype 1, a heterozygous structural variant on haplotype 2, or a homozygous structural variant affecting both haplotypes. If positions *i* and *j* are on the same phase block *m*_*i*_ = *m*_*j*_, then the haplotypes of *i* and *j* must match. Thus, we calculate the log-likelihood ratio 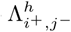 to be the maximum of 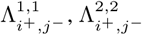, and *Λ*_*i*^+^,*j*^‒^_, corresponding to a novel adjacency resulting from a heterozygous structural variant on haplotype 1, a heterozygous structural variant on haplotype 2, or a homozygous structural variant,

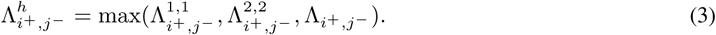

However, if positions *i* and *j* are not on the same phase block, *m*_*i*_ ≠ *m*_*j*_, then the haplotypes of *i* and *j* may not match. For example, SNPs assigned to haplotype 1 of phase block *m*_*i*_ may originate from the same chromosome as SNPs assigned haplotype 2 of phase block *m*_*j*_. In this case, we calculate the log-likelihood ratio 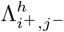 as,

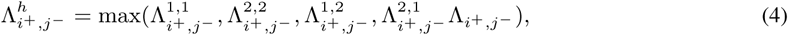

which accounts for the ambiguity in haplotype assignment. NAIBR reports both the phased log-likelihood ratio 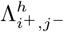 as well as the inferred haplotype for each novel adjacency.

## 3 Results

We assess NAIBR’s ability to detect novel adjacencies in simulated and real 10X long-read sequencing data and benchmark against 5 other methods: Long Ranger (Zheng *et al*., 2016), GROC-SVs (Spies *et al*., 2016), GASV (Sindi *et al*., 2009b), GASVPro (Sindi *et al*., 2012), and LUMPY (Layer *et al*., 2014). We chose these methods for comparison because they utilize different combinations of signals to identify and rank novel adjacencies. Long Ranger is 10X Genomics’ structural variant detection program. Long Ranger identifies novel adjacencies by computing overlapping pairs of linked-reads and computes a likelihood score based on the number of overlaps observed in the data. GROC-SVs is also designed for linked-read sequencing data. GROC-SVs identifies structural variants by performing local assembly on barcoded reads. GROC-SVs operates in two steps. First it assigns p-value to novel adjacencies, and then it performs local assembly, labelling novel adjacencies as assembled or unassembled. In some cases, unassembled variants have smaller p-values than assembled variants. GASV, GASVPro, and LUMPY analyze paired-end sequencing data. When running these algorithms on 10X Genomics data we ignore barcodes and treat the data as Illumina paired-end sequencing data. GASV uses only discordant read-pairs and ranks novel adjacencies by the number of supporting discordant read-pairs. GASVPro uses a combination of discordant read-pairs and breakend read depth to assign a log-likelihood score to each predicted novel adjacency. LUMPY uses discordant and split reads to call novel adjacencies and reports a p-value for each adjacency.

We benchmark NAIBR against each method on both simulated and real data. Reported novel adjacencies are ranked from highest to lowest confidence according to the metrics used by each method. We run each method using its default parameters. More specifics can be found in the Supplement.

We simulate several types of structural variants – including duplications, deletions, translocations, and inversions – on chromosomes 17 and 18 of the human reference genome hg19. To assess NAIBR’s ability to detect novel adjacencies that occur on a single haplotype, we simulate two test genomes, one that contains 400 homozygous structural variants and one that contains 400 different structural variants on each haplotype. Translocations and inversions create more than 1 novel adjacency in the simulated genome, resulting in 508 homozygous novel adjacencies in the first simulated genome and 1027 heterozygous novel adjacencies in the second simulated genome. We simulate linked-read sequencing to 30X coverage. Details on simulation can be found in the Supplement. Figure 4a shows the precision-recall curve for all 5 methods run on the 30X test dataset containing 508 homozygous novel adjacencies. NAIBR has the highest recall of all methods, correctly identifying 479/508 homozygous novel adjacencies. GASV correctly identified 309/508 variants, however GASV reported several thousand variants, resulting in very low precision at high values of recall. LUMPY performed similarly to GASV, correctly identifying 308/508 true variants, however the algorithm only reports variants with high probability scores according to their scoring metric, resulting in lower recall. GASVPro identified as many true variants as NAIBR at 50% recall, but only reported 289/508 true variants in total, compared to 479 reported by NAIBR.

**Figure 4:**
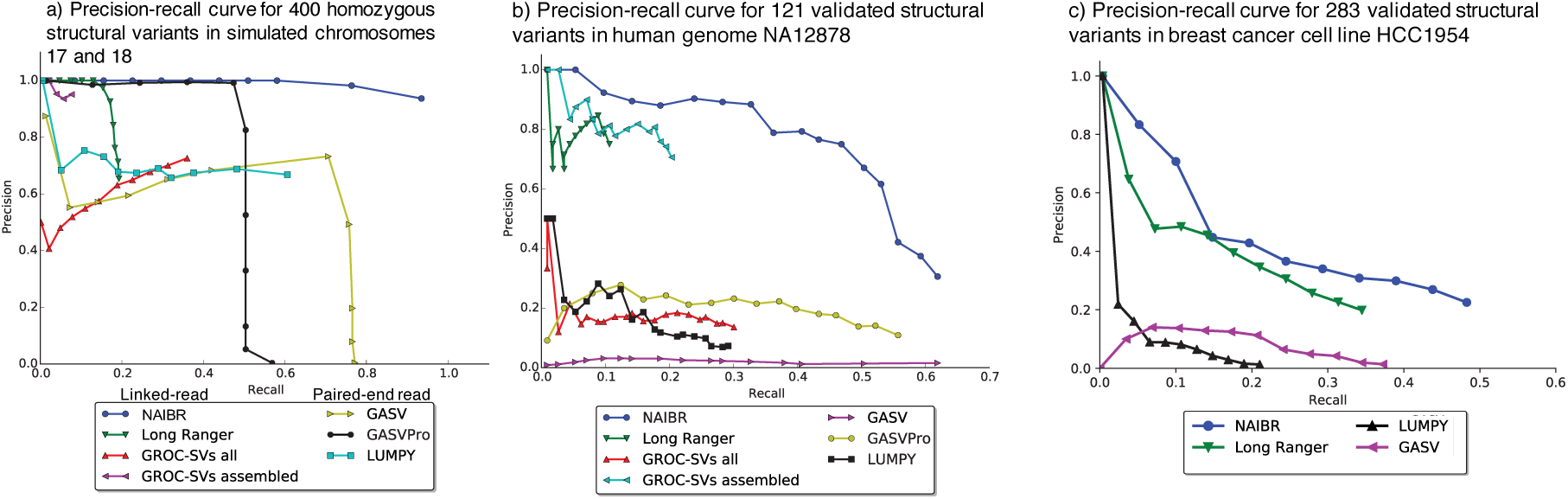
**a)** Precision-recall curve for NAIBR, Long Ranger, GROC-SVs, GASV, GASVPro, and LUMPY on 30X simulated data from chromosomes 17 and 18, containing 400 homozygous structural variants. b) Precision-recall curve for NAIBR, GASV, GASVPro, LUMPY, GROC-SVs, and Long Ranger evaluated against the set of 123 validated structural variants > 7Kbp from NA12878 (Layer *et al*., 2014). c) Precision-recall curve for NAIBR, GASV, GASVPro, LUMPY, GROC-SVs, and Long Ranger evaluated against the set of validated structural variants ≥ 30Kbp from breast cancer cell line HCC1954 (Bignell *et al*., 2007; Stephens *et al*., 2009; Galante *et al*., 2011).

Long Ranger and GROC-SVs are each designed to utilize linked-reads, however both methods are limited to the identification of certain types of variants. Long Ranger reports variants larger than 30Kbp and GROC-SVs only reports variants larger than 10Kbp. The simulated dataset contains 369 novel adjacencies larger than 10Kbp and 269 structural variants larger than 30Kbp. Both Long Ranger and GROC-SVs perform with lower precision than NAIBR. For GROC-SVs, 38/39 of assembled novel adjacencies were present in the truth set, showing that the local assembly approach has high precision. However an additional 59 true novel adjacencies were predicted by GROC-SVs but failed to assemble, indicating that local assembly removes many true positives. We perform the same comparison on a simulated dataset containing 800 heterozygous novel adjacencies with similar results. We also compare the runtime and memory usage of NAIBR to other methods and find that NAIBR outperforms other linked-read methods (see Supplement). These results demonstrate that NAIBR’s incorporation of both linked-read and paired-end read data improves performance over other methods without significant additional time or memory requirements.

### 3.1 Benchmarking on NA12878

To assess NAIBR’s ability to detect known variants from a real dataset, we obtained whole-genome sequencing data of individual NA12878of the 1000 Genomes Project from 10X genomics (https://support.10xgenomics.com/genome-exome/datasets/NA12878_WGS_210). The data was sequenced on the Chromium platform to 35X sequencing coverage with the Illumina Hiseq2500 to produce 845 million 98bp reads with mean insert size of 340bp and a mean molecule size of 68Kbp. We used the 2,950 validated novel adjacencies in NA12878 reported in Layer *et al*. (2014) as the truth set. Novel adjacencies in this dataset were validated by split-read mapping analysis of independent long-read sequencing data from PacBio or Illumina Moleculo platforms.

Figure 4b shows the precision-recall curves each method against 121 validated structural variants larger than 7Kbp. NAIBR correctly predicted 73/121 novel adjacencies > 7Kbp. GASV also correctly predicted 73 novel adjacencies but displayed poor precision. GROC-SVs reported variants that it was able to assemble using its local assembly pipeline as well as variants that were not assembled. The local assembly step of GROC-SVs drastically reduces the number of reported false positives but also fails to assemble several predictions matching the truth set. Long Ranger reported 17 novel adjacencies, correctly identifying 12/32 variants larger than 30Kbp from the truth set, compared to 17/32 variants larger than 30Kbp detected by NAIBR. Figure S9 shows the precision-recall curves for each of the 5 methods on different structural variant sizes, ranging from 50bp-30Kbp. NAIBR outperforms other linked-read methods at detecting variants between 50bp and 10Kbp and outperforms paired-end methods at detecting variants larger than 5Kbp. NAIBR outperforms both linked-read and paired-end methods at detecting variants between 1Kbp and 10Kbp. In summary, NAIBR detects large structural variants (> 1Kbp) with better precision than paired-end callers GASV, GASVPro, and LUMPY and has higher recall and precision for variants smaller than 10Kbp compared to linked-read callers Long Ranger and GROC-SVs.

### 3.2 Tumor cell line HCC1954

We test NAIBR’s ability to detect somatic structural variants in tumor cell line HCC1954T. The cell line was derived from a grade 3 invasive ductal carcinoma and sequenced by 10X Genomics to 35X coverage with a mean molecule size of 85Kbp. The matched normal HCC1954N was sequenced by 10X Genomics to 35X coverage with a mean molecule size of 88Kbp. We identify novel adjacencies in both HCC1954T and HCC1954N using 4 different methods: NAIBR, Long Ranger, GASV, and LUMPY. We formed a set of 369 true novel adjacencies by combining PCR-validated novel adjacencies from three previous studies: Bignell *et al*. (2007), Stephens *et al*. (2009), and Galante *et al*. (2011).

NAIBR identifies 142 PCR-validated novel adjacencies, significantly more than Long Ranger (100), GASV (117), and LUMPY (55) (Figure S6). NAIBR also demonstrates better precision at all levels of recall than other methods (Figure 4c). Notably, GASV has significantly lower precision than Long Ranger and NAIBR, predicting over ten times as many novel adjacencies with lower recall than the other methods (Figure S6). NAIBR significantly outperforms Long Ranger at identifying duplications and interchromosomal events (Figure S5), identifying 72 duplications compared to 50 identified by Long Ranger and 53 interchromosomal events compared to 37 predicted by Long Ranger. Over 30% of the breakends predicted by NAIBR lie within 50bp of the breakends of the PCR-validated novel adjacencies, compared to approximately 10% of breakends predicted by Long Ranger and GASV (Figure S7).

Several of the novel adjacencies in HCC1954T that were not identified by Long Ranger and GASV affect known oncogenes and tumor suppressors. For example, a novel adjacency between Chr11:93153935 and Chr11:93160223 occurs within the gene CCDC67, potentially leading to loss of function of the gene. CCDC67 has recently been identified as a tumor suppressor gene (Yin *et al*., 2016; Zhu *et al*., 2014). A novel adjacency between Chr14:89829140 and Chr7:155683934 affects the forkhead transcription factor checkpoint suppressor, CHES1. CHES1 expression has been shown to be reduced across many cancer types (Huot *et al*., 2014). A novel adjacency between Chr11:69059340 and Chr11:69089741, surrounding the MYEOV gene (Chr:69061613-69064754), is potentially a result of a duplication of MYEOV. MYEOV has been shown to be amplified in breast cancer patients and has been identified as a candidate oncogene (Janssen *et al*., 2002).

## 4 Discussion

We present NAIBR, a probabilistic algorithm for the identification of novel adjacencies using linked-read sequencing data. Linked-read sequencing combines low per-base error rate of short-read sequencing technologies with long-range linking information of long-read technologies. Linked-read sequencing offers drastically improved mapping and phasing results compared to paired-end sequencing (Bishara *et al*., 2015) with similar cost, making it an attractive option for researchers. NAIBR is one of the first algorithms that identifies structural variation by using signals unique to linked-read sequencing data. NAIBR uses discordant read-pairs obtained from paired-end reads combined with candidate split molecule obtained from linked-reads to identify and rank novel adjacencies. NAIBR detects novel adjacencies with higher accuracy and precision than existing methods on both simulated and real linked-read sequencing data.

We also demonstrate NAIBR’s ability to predict somatic novel adjacencies from cancer data by applying it to cell line HCC1954. Several novel adjacencies detected by NAIBR were not identified by other methods, including novel adjacencies affecting tumor suppressor genes CCDC67 and CHES1 and candidate oncogene MYEOV. While some of the novel adjacencies predicted by NAIBR might be verified PCR, it is possible that long-read sequencing data (e.g.from PacBio or Oxford Nanopore) would be required, since these adjacencies were not readily apparent from short-read sequencing data.

As future work, we plan to incorporat.epse additional signals, such as read depth and split-reads, into our probabilistic model for identifying novel adjacencies. Our algorithm can also be extended by performing local assembly on linked-reads supporting novel adjacencies to reconstruct structural variants, as is done by GROC-SVs (Spies *et al*., 2016).

The utility of linked-reads in identifying novel adjacencies between nearby positions on the reference genome (such as a small deletion) is limited by the fact that each molecule is sequenced to low coverage, introducing large gaps between read-pairs sequenced from the same molecule. Thus, with the current Chromium technology from 10X Genomics, linked-reads provide substantial additional power to detect large structural variants but provide little additional power above that of paired-end sequencing in the detection of small structural variants. However, linked-read sequencing is a very new technology and is likely to improve in the coming years. As molecules are sequenced to higher coverage, NAIBR’s ability to detect novel adjacencies using candidate split molecules will improve significantly, enabling the identification of novel adjacencies arising from smaller structural variants.

## NAIBR Software

Software is available at compbio.cs.brown.edu/software

